# A Massively Parallel Trafficking Assay Accurately Predicts Loss of Channel Function in *KCNH2* Variants

**DOI:** 10.1101/2021.07.10.451881

**Authors:** Chai-Ann Ng, Rizwan Ullah, Jessica Farr, Adam P. Hill, Krystian A. Kozek, Loren R. Vanags, Devyn Mitchell, Brett M. Kroncke, Jamie I. Vandenberg

## Abstract

High throughput genomics has greatly facilitated identification of genetic variants. However, determining which variants contribute to disease causation is challenging with more than half of all missense variants now classified as *variants of uncertain significance* (VUS). A VUS leaves patients and their clinicians unable to utilize the variant information in clinical decision-making. In long QT syndrome type 2, *KCNH2* channel function is directly associated with disease presentation. Therefore, functional phenotyping *of KCNH2* variants can provide direct evidence to aid variant classification. Here, we investigated the expression of all codon variants in exon 2 of *KCNH2* using a massively parallel trafficking assay and for a subset of 458 single nucleotide variants compared the results with peak tail current density and gating using automated patch clamp electrophysiology. Trafficking could correctly classify loss of peak tail current density variants with an AUC reaching 0.94 compared to AUCs of 0.75 to 0.8 for *in silico* variant classifiers. We suggest massively parallel trafficking assays can provide prospective and accurate functional assessment for all missense variants in *KCNH2* and most likely many other ion channels and membrane proteins.

## Introduction

Identifying the specific genetic cause in inherited disorders can be particularly valuable for patient management as well as cascade screening for family members (Semsarian et al., 2021). However, most variants are ultra-rare and therefore compelling clinical data to establish whether a variant identified in a gene of interest is the cause of disease is often absent. For many disease genes the majority of variants are currently classified as variants of uncertain significance (Landrum et al., 2018; Lek et al., 2016), leaving patients, their families and doctors in genetic limbo. Overcoming this hurdle is critical to realising the potential of genomic medicine.

Long QT Syndrome (LQTS) is a myocardial repolarization disorder that predisposes affected individuals to *Torsades de Pointes*, a life-threatening ventricular arrhythmia (Priori et al., 2013). Around 75 % of all LQTS cases result from mutations in three cardiac ion channel genes; *KCNQ1* (coding K_V_7.1 channel α-subunit, conducting *I*_Ks_; LQTS type 1), *KCNH2* (a.k.a. hERG, coding K_V_11.1 channel α-subunit, conducting *I*_Kr_; LQTS type 2), and *SCN5A* (coding Na_V_1.5 channel α-subunit, conducting *I*_Na_; LQTS type 3) (January et al., 2000; Keating & Sanguinetti, 2001; Kroncke et al., 2018; Rajamani et al., 2002; Roden & Balser, 1999). All three genes are included in the American College of Medical Genetics and Genomics list of medically actionable genes due to their association with increased risk of sudden cardiac death in otherwise healthy individuals (Kalia et al., 2017). Assigning risk of LQTS for carriers of these variants is of particular interest as effective therapy (beta-blockers) is available (Priori et al., 2013) and it greatly facilitates cascade screening of family members.

In contrast to characterizing variants discovered in clinically-identified probands and their families, *in silico* and *in vitro* experiments may *prospectively* identify variants that cause channel dysfunction before they are observed in individuals. Previously, characterization of variants in cardiac ion channels has relied on manual patch clamp electrophysiology, which is prohibitively time consuming for thousands of variants. Two recent technological developments have enabled the functional characterisation of variants at a scale commensurate with their rate of discovery: an automated patch clamp platform, which permits analysis of hundreds of cells simultaneously (Glazer, Wada, et al., 2020; Ng et al., 2020; Vanoye et al., 2018); and massively parallel variant effect mapping coupled to cell survival (Findlay et al., 2018) or expression/trafficking assays (Kozek et al., 2020). The automated patch clamp platform enables phenotyping of current density, gating, and permeation properties of variant channels, both homozygous and heterozygous (Ng et al., 2020), at moderate throughput (hundreds of variants per month). Massively parallel variant effect mapping enables the analysis of tens of thousands of variants but at a reduced depth of information (Glazer, Kroncke, et al., 2020; Kozek et al., 2020). For *KCNH2*, massively parallel expression/trafficking assays (Kozek et al., 2020) have great potential as the deleterious effects of ion channel variants in *KCNH2* are largely due to trafficking defects (Anderson et al., 2014; Delisle et al., 2004).

The aim of this study was to assess the feasibility of prospectively characterizing all *KCNH2* missense variants. We investigated all missense single nucleotide variants (SNVs) in Exon 2 of *KCNH2* and compared results from patch clamp electrophysiology, the benchmark in ion channel characterization, with a massively parallel variant effect map of channel trafficking. Exon 2, which encodes the first 77 residues of the PAS domain, is a known hotspot for pathogenic variants (Shimizu et al., 2009). In total, 458 non-redundant, missense SNVs were compared between the two methods. Forty-one percent of variants had at least 50 % trafficking defect and nearly 90 % of these had a dominant negative phenotype based on heterozygous peak tail current density measurements. Only 4.5 % of variants showed a less severe phenotype when assessed using heterozygous peak tail current density measurements. There was a strong correlation between homozygous trafficking defects and heterozygous peak tail current density (Spearman rank-order correlation ρ = 0.75, p<0.001; Table 1), higher than the moderate correlation between heterozygous peak tail current density and the bioinformatic prediction algorithms, REVEL (Ioannidis et al., 2016) or CardioBoost (Zhang et al., 2021).

**Table 1.** Spearman rank-order correlations between peak tail current density and predictive features including trafficking. All correlations significant with a nominal p-value < 0.001.

| FEATURE | SPEARMAN $\rho$ [95% CI] |
| --- | --- |
| REVEL | 0.52 [0.46 – 0.58] |
| CARDIOBOOST | 0.52 [0.45 – 0.58] |
| STRUCTURE-DERIVED<br>FEATURE | 0.42 [0.34 – 0.50] |
| TRAFFICKING | 0.75 [0.70 – 0.80] |

This is the first study to independently validate large-scale variant pathogenicity of ion channel variants measured by a massively parallel trafficking assay with detailed electrophysiology data. We suggest the assays described here will enable the prospective collection of meaningful variant data for all missense variants in *KCNH2*. Furthermore, similar massively parallel trafficking assays will be useful for any ion channels and membrane proteins where protein misfolding is the major cause of loss-of-function (Yue et al., 2005). Resources such as these will become increasingly important to address the deluge of variants of uncertain significance arising from the increasing prevalence of genome sequencing in affected and unaffected populations (Starita et al., 2017).

## Results

### Massively parallel trafficking assay

For the present study, we selected Exon 2, a “hotspot” for known pathogenic variations (Shimizu et al., 2009; Vandenberg et al., 2012). We followed a previously published protocol to generate a variant effect map for K_V_11.1 (*KCNH2*) trafficking (Kozek et al., 2020). A total of 1,463 missense out of a possible 1,540 variants (95 %) and 77 nonsense variants were generated. Trafficking scores were calculated by counting variants in pools of sorted cells as previously described (Kozek et al., 2020) and detailed in the materials and methods. Final trafficking scores were calculated using at least three replicates from two separate variant-barcode plasmid constructs. The trafficking scores ranged between 0 and 160 % of WT for the 1,463 missense variants identified, 0-165 % of WT for the 77 synonymous variants (mean of 95 and median of 100 % of WT) and 0 % of WT for the 77 nonsense variants (mean and median of 0 % of WT; Figure 1). The distribution of the trafficking scores was bimodal with peaks around 0 and 100, suggesting most variants are well described as loss-of-function/deleterious or no-change/benign (Figure 1B). A summary heatmap of the trafficking scores, categorized into five groups (0-25 %, 25-50 %, 50-75 %, 75-150 % and >150 %), is shown in Figure 1C. Interestingly, several residues tolerate only substitutions of amino acids with similar chemical properties to WT, e.g., I30 and I31 only tolerate branched aliphatic residue substitutions while Y43 and Y54 only tolerate aromatic residue substitutions. These constraints likely indicate the nature of interactions of these residues in the folded protein.

**Figure 1:**
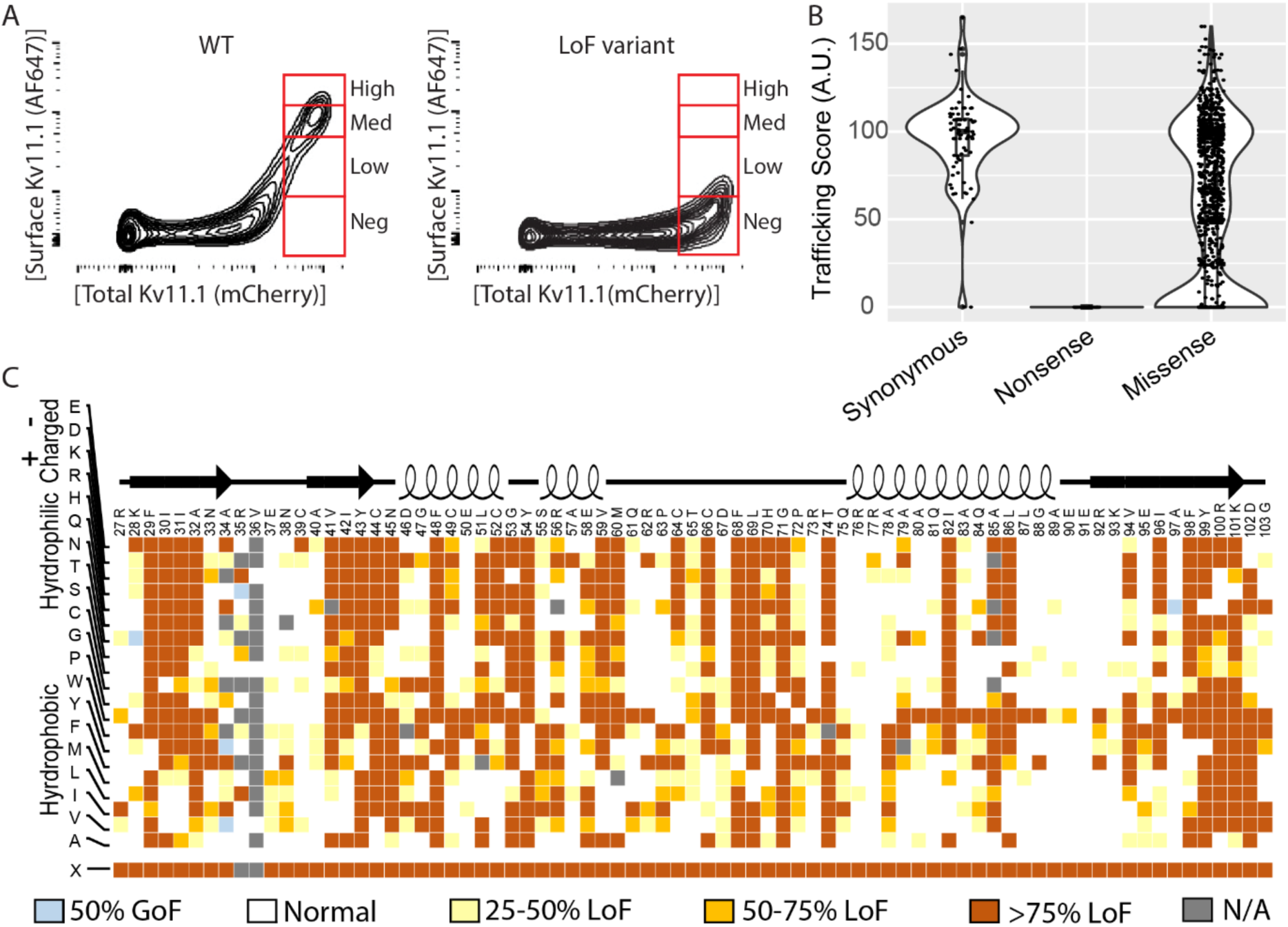
Deep mutational scan of exon 2 of *KCNH2* identifying trafficking deficient variants. (A) Trafficking pattern of wild type (WT) *KCNH2* compared to loss of function (LoF) variants based on the expression of Alexa Fluor 647 (AF647) denoting the surface expression of K_V_11.1. X-axis is mCherry fluorescence (indicating K_V_11.1 expression), the y-axis is surface stained K_V_11.1 (protein product of *KCNH2*). (B) Violin plots showing the trafficking scores for all the synonymous, nonsense and missense variants in the exon 2 region. (C) A total of 1,540 variants were observed and are colour coded based on their trafficking patterns. The different levels of trafficking are normalized to WT (synonymous) and categorized as: greater than 75 % LoF (brown; 554 variants), 50-75 % LoF (orange; 100 variants), 25-50 % LoF (yellow; 201 variants), WT-like (white; 642 variants) and GoF (blue; 5 variants).

### Large-scale automated patch clamp assay

Most variants observed clinically involve single nucleotide changes. For our automated patch clamp assays of heterozygous *KCNH2* variants, we therefore characterized the subset of 458 non-redundant missense SNV in exon 2. Typical examples of tail currents from a selection of 11 variants stably expressed in Flp-In HEK293 cell lines (and a negative control) recorded on the same 384-well plate are shown in Figure 2A. The current densities recorded at −120 mV are presented as violin plots in Figure 2B. The median values for all 458 variants, categorized into five groups (0-25 %, 25-50 %, 50-75 %, 75-150 % and >150 % of WT levels), are shown in Figure 2C.

**Figure 2:**
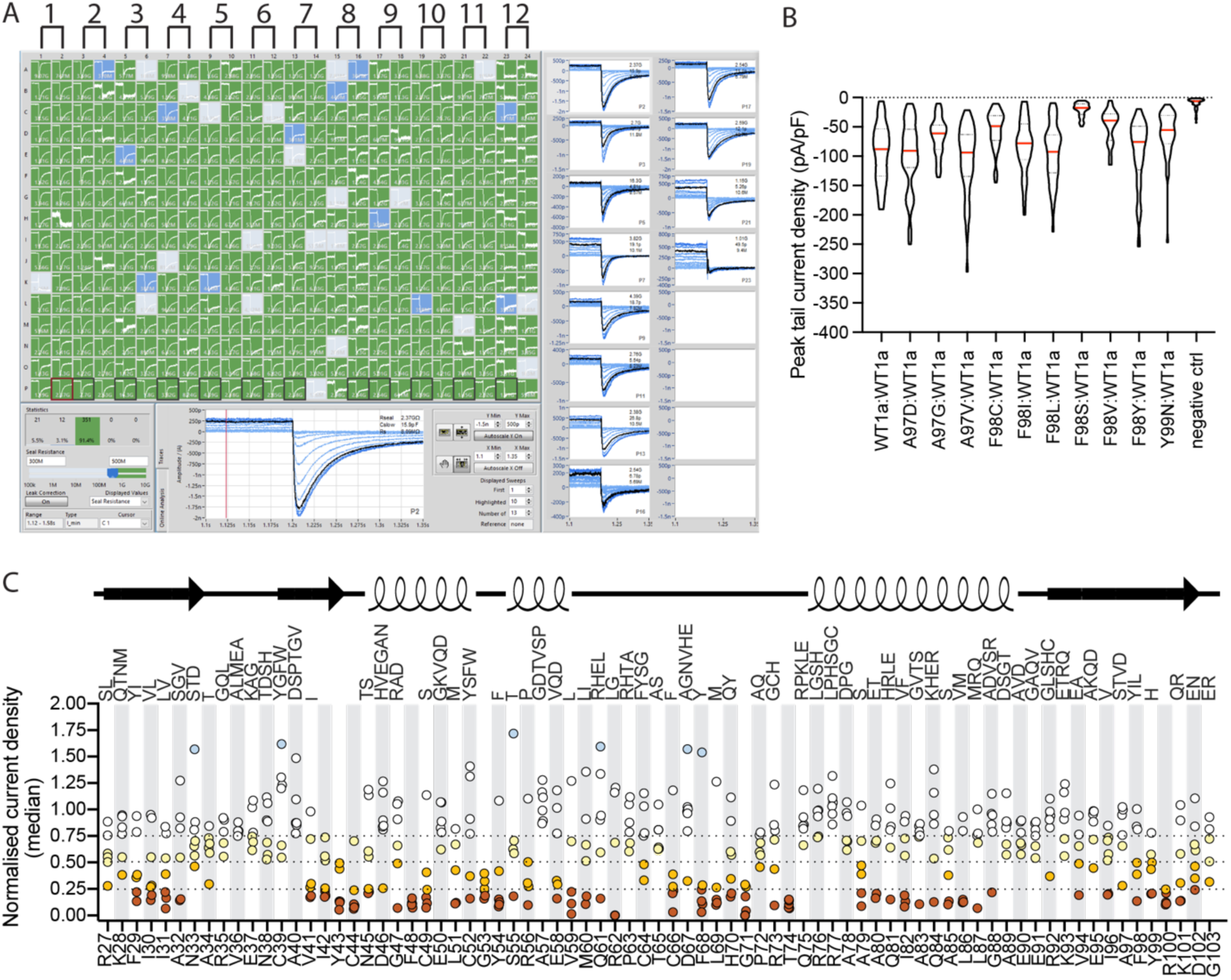
Functional phenotyping of all the missense SNV in the exon 2 of *KCNH2* using Automated Patch-Clamp (APC). (A) Patch Control software showing a range of automatically scaled peak tail current traces correspond to the WT:WT (1), A97D:WT (2), A97G:WT (3), A97V:WT (4), F98C:WT (5), F98I:WT (6), F98L:WT (7), F98S:WT (8), F98V:WT (9), F98Y:WT (10), Y99N:WT (11) and an negative control (12). The highlighted black peak tail currents were acquired after depolarization at +40 mV for 1-s before stepping to –120 mV. More than 90 % of patch-clamp recordings (green wells) have seal resistance greater than 500 M*Ω*. A total of 64 patch-clamp recordings (32×2) will be collected for each of these variants in technical replicate. (B) The violin plots for the –120 mV peak tail current density that correspond to those variants shown in panel A. (C) The median current density of all the missense single nucleotide variants normalized to the median current density of WT. The n number used to derive these median values are from a total of 22,850 patch-clamp experiments. The different levels of peak tail current density are categorized as: greater than 75 % LoF (brown; 109 variants), 50-75 % LoF (orange; 55 variants), 25-50 % LoF (yellow; 93 variants), WT-like (white; 195 variants) and GoF (blue; 6 variants). 72/141, 47/118 and 82/199 variants located within the *α*-helices, *β*-sheets or loop region have WT-like peak tail current density (i.e. >0.75), respectively.

For each individual patch-clamp recording that had sufficient peak tail current density (see methods), we assessed channel gating. Example data sets for the variants with the most perturbing effect on steady-state activation (M60V), inactivation (E90K) and deactivation (N33D) are shown in Figure 3A (panels i, ii, iii, respectively). Summary results for these three variants are shown as violin plot insets in Figure 3A. No variants resulted in more than 10 mV shift in the midpoint of steady-state activation (Supplementary Figure 1A). Steady-state inactivation was assessed by quantifying the ratio of peak tail current at −50 mV and −120 mV. Fifteen variants have enhanced inactivation (25-50 % less peak tail current at −50 mV compared to WT, yellow squares in Figure 3B, top panel) and four variants had greater current at −50 mV, blue squares in Figure 3B, top panel). None of these variants had significant effects on the rates of inactivation (Supplementary Figure 1B). Twenty-one variants accelerate channel deactivation, assessed as the ratio of current decay after 500 ms at −50 mV while 4 variants slowed channel deactivation (Figure 3B, bottom panel). When assessing deactivation by calculating a weighted *τ* for the current decay at −50 mV there were 36 variants that had a reduced weighted *τ* value more than 25 % from WT level (Supplementary Figure 1C).

**Figure 3:**
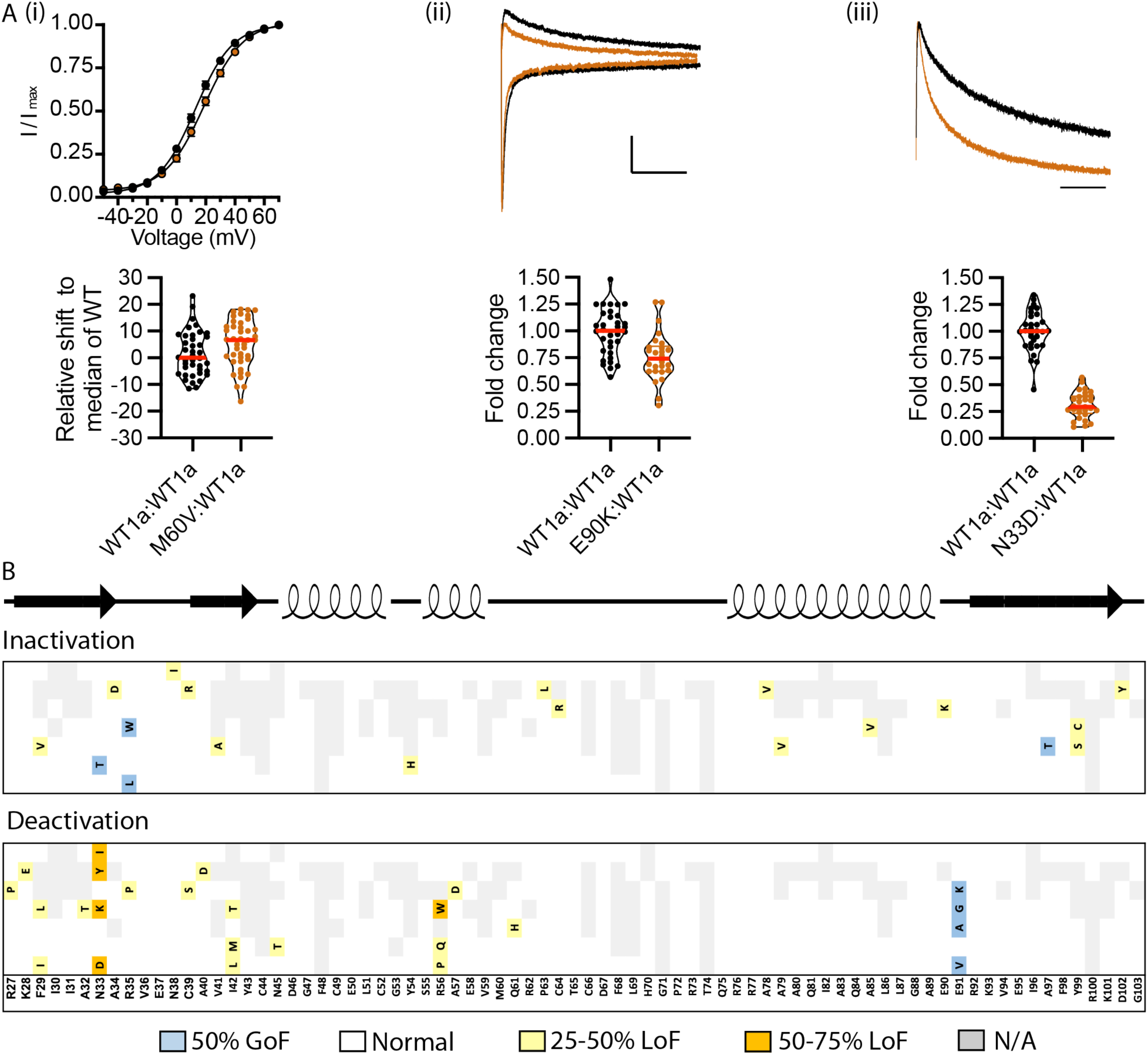
Gating phenotypes of all the missense SNV in the exon 2 of *KCNH2*. (A) Example of variants that have the most perturbed effect for (i) *V*_0.5_ of activation determined by fitting a Boltzmann function to the –120 mV tail current after 1-s of depolarization and the relative shift from the median of WT; (ii) Channel inactivation by taking the ratio for the –50 and –120 mV current amplitudes; (iii) Channel deactivation by determining the decay of the current at –50mV after 500 ms (current amplitude_500ms_/current amplitude_peak_). (B) Summary heatmaps for all the missense single nucleotide variants that have sufficient current density to be reliably analyzed. All the hyperpolarized shift and less than +10 mV depolarized shift in *V*_0.5_ from WT are categorized as WT-like (white). For channel inactivation and onset of channel deactivation, variants are colour coded as having greater than 75 % LoF (brown), 50-75 % LoF (orange), 25-50 % LoF (yellow), WT-like (white) and GoF (blue). When the n number that pass quality control for analysis of gating parameters is less than 25 % of the recordings that have passed quality control parameters for peak tail current density then these variants are classified as not analyzed (N/A).

### Trafficking-defects are the dominant loss-of-function phenotype

Though the expression systems differed between the trafficking (homozygously expressed) and automated patch clamp (heterozygously expressed) assays, trafficking and peak tail current density results were similar (Figure 4A-B); Spearman ρ = 0.75 [95 % confidence interval: 0.70-0.80] (Table 1). Overall, 41 % of variants trafficked less than 50 % of WT while 36 % of variants had less than 50 % peak tail current density compared to WT. Similarly, the distribution of high impact variants which affect trafficking and peak tail current density map to the same clusters in the K_V_11.1 structure (Figure 4C). Residues with the greatest tolerance to substitution reside peripherally in the domain, typically facing the aqueous, intracellular environment. Residues intolerant to substitution were largely in the interior of the domain. There were only 15 homozygous variants with a severe trafficking defect (< 15 % of WT) but a heterozygous peak tail current density between 40-60 % of WT (Figure 5B). This suggests that nearly 90 % of trafficking defective variants are dominant negative. At the other extreme—trafficking defective variants apparently rescued by co-expression with the WT allele—there were only 21 variants which trafficked 20-60 % of WT yet produced a peak tail current density >75 % of WT (Figure 5C). These results indicate that most channel variants, though not all, are well described in a homozygous expression context.

**Figure 4:**
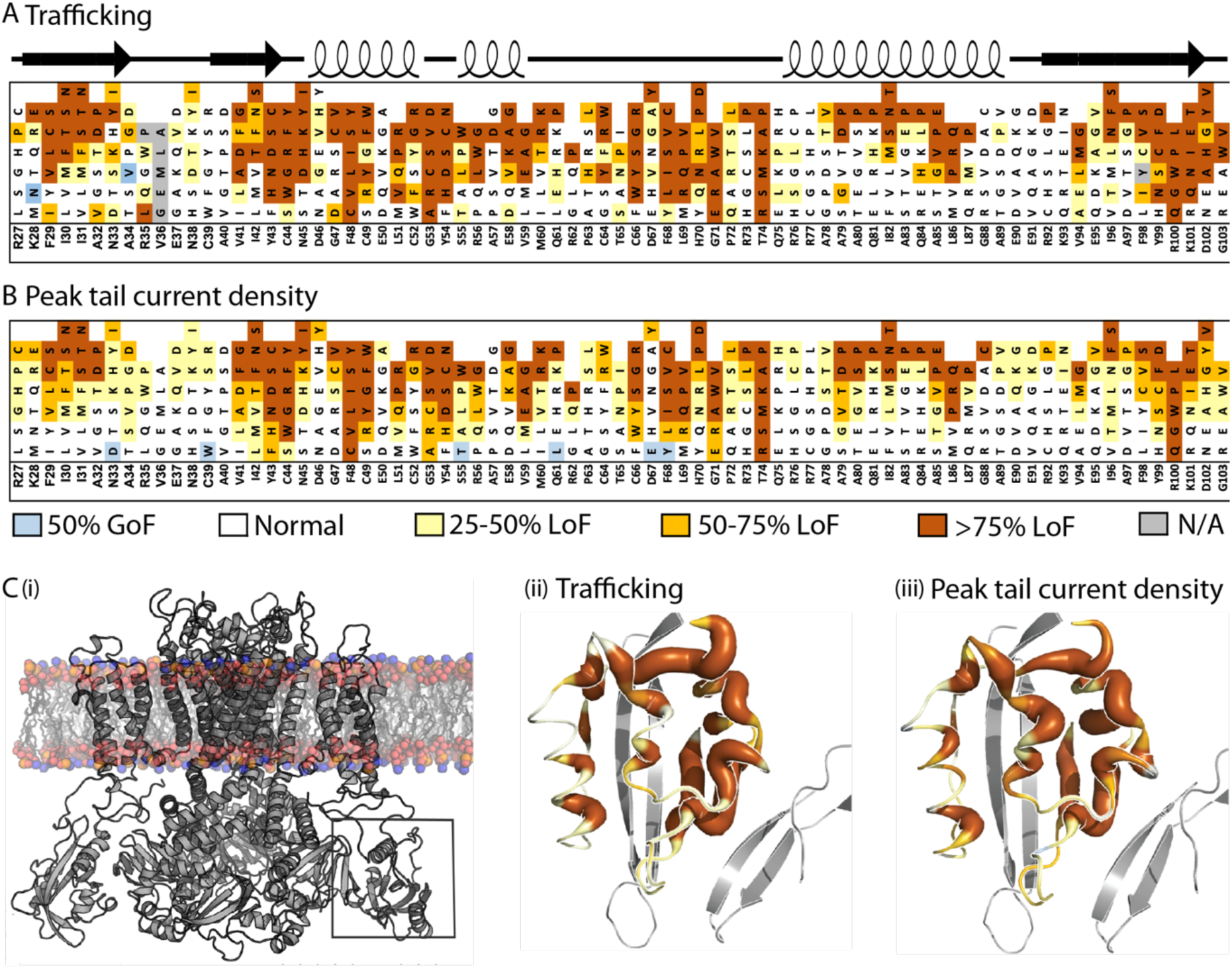
One third of all the missense SNV in the exon 2 of *KCNH2* cause at least 50 % loss of function. (A) Expression defects for homozygous missense SNV and (B) peak tail current density for heterozygous missense SNV are colour coded as having greater than 75 % LoF (brown), 50-75 % LoF (orange), 25-50 % LoF (yellow), WT-like (white) and GoF (blue). (C) Cryo-EM structure of K_V_11.1 K^+^ ion channel (PDB ID: 5VA3) embedded in a lipid bilayer with the location of the exon 2 residues (amino acids 27-103) highlighted within a box (i-ii). Mapping of trafficking (ii) and peak tail current density (iii) to the structure of the PAS domain. Thicker representation, in addition to colour, indicates a more deleterious phenotype.

**Figure 5:**
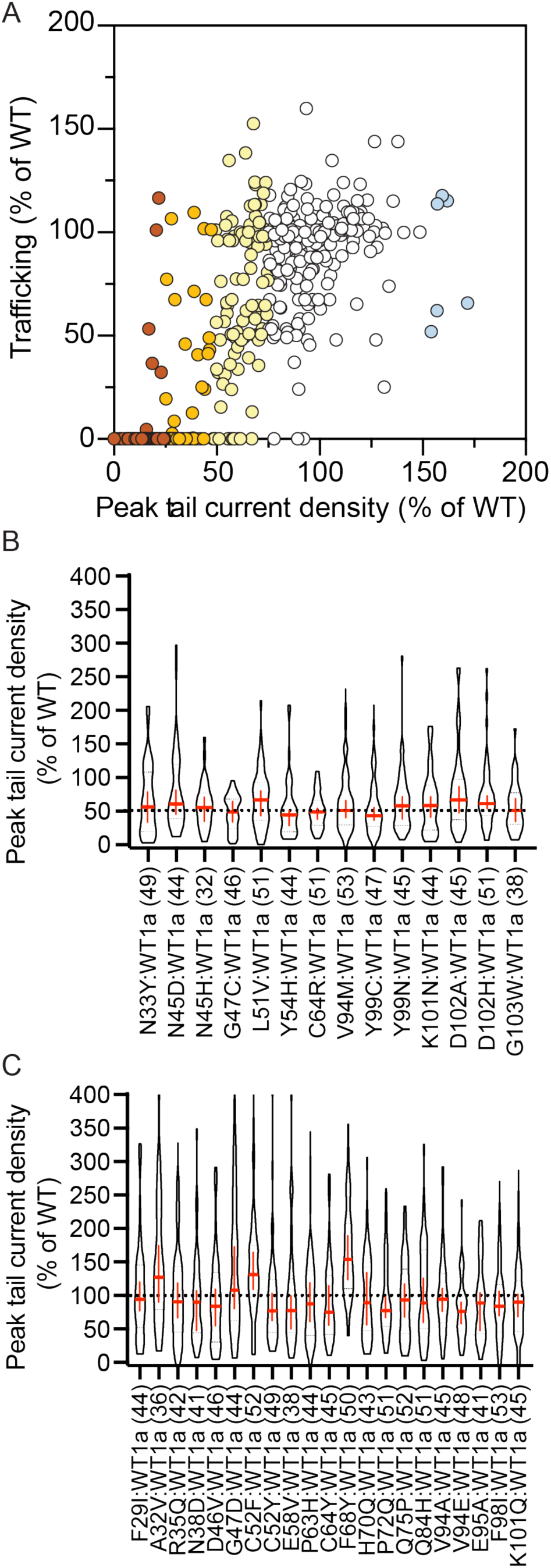
Haploinsufficient *KCNH2* variants. (A) Correlation between the homozygous trafficking to heterozygous peak tail current density (Spearman ρ = 0.75). These variants are colour coded based on the peak tail current density as having greater than 75% LoF (brown), 50-75% LoF (orange), 25-50% LoF (yellow), WT-like (white) and GoF (blue). (B) Haploinsufficient variants identified by trafficking less than 15 % of WT and peak tail current density between 40 and 75 % in panel A. (C) Variants that can be rescued by co-expressed with WT (trafficking <60 % of WT and peak tail current density >75 % of WT in panel A). Red crosses within the violin plot are median with 95 % confidence interval and n numbers for each variant are in the bracket.

In addition, few variants reduced channel function solely through defective gating. Most channel gating defects were observed concomitant with significantly disrupted trafficking and peak tail current density. Notably, this was the case for residues located within the hydrophobic patch on the surface of the PAS domain (e.g. K28E, F29L, I31S, I42N and Y43C) previously shown to be important for binding the PAS domain to the cNBH domain (Ke et al., 2013; Morais Cabral et al., 1998) with one notable exception, N33D (see Figure 4A-B).

### A comparison between bioinformatic predictions and functional data

To assess the concordance between these tools and our functional data, we compared our *in vitro* results with the Rare Exome Variant Ensemble Learner (REVEL) (Ioannidis et al., 2016) and CardioBoost (Zhang et al., 2021) *in silico* variant classifiers. REVEL aggregates 13 tools to compute a score between 0 and 1 (Figure 6A). CardioBoost is a recent method optimized specifically to predict pathogenicity of variants in inherited heart disease-associated genes (Figure 6B). We additionally evaluated a structure-derived feature which measures of the density of pathogenic variants in three-dimensional space (previously referred to as “functional density” (Kroncke et al., 2019), here called “structure-derived feature” to avoid confusion with peak tail current density; Figure 6C). We found a moderate correlation between REVEL and CardioBoost and peak tail current density, both with a Spearman *ρ* = 0.52, and 66 false-positive variants. CardioBoost produced more false negatives (35) compared to REVEL (16) when compared to the peak tail current density measurements (Figure 6). The structure-derived feature performed slightly worse than either REVEL or CardioBoost with a Spearman *ρ* = 0.42.

**Figure 6:**
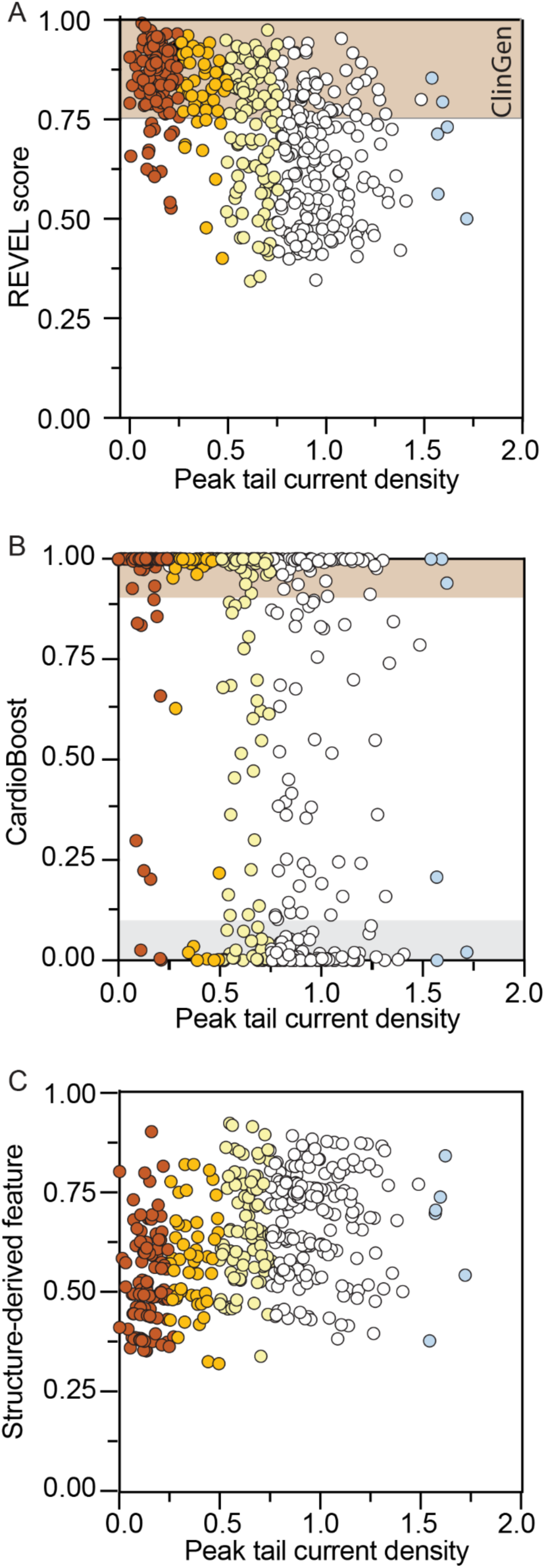
Bioinformatic prediction overcalling the loss-of-function *KCNH2* variants. (A) Correlation between the heterozygous peak tail current density to REVEL score (Spearman ρ = 0.52). Sixty-six variants that have WT-like peak tail current density were overcalled by REVEL score as LoF when using the more stringent Clin-Gen cut-off of 0.75 (Rehm et al., 2015). Only two variants that have less than 50 % WT peak tail current are called as normal by REVEL score of <0.5. (B) Correlation between the heterozygous peak tail current density to CardioBoost (Spearman ρ = 0.52). Sixty-six variants that have WT-like peak tail current density were overcalled by CardioBoost as disease-causing (*≥* 0.9). Only nine variants with less than 50 % WT peak tail current density were assigned benign/likely benign (*≤*0.1). (C) Correlation between the heterozygous peak tail current density to structure-derived feature (Spearman ρ = 0.42).

### Massively parallel trafficking assay predicts loss-of-function *KCNH2* variants

To assess the overlap of information between *in silico* predictions and the trafficking data we generated, we built predictive models of peak tail current density using *in silico* (REVEL, CardioBoost, and the structure-derived feature) predictors and trafficking data as features (Figure 7). The coefficient of determination, adjusted for the number of regression features, predicting peak tail current density using *in silico* features alone was R^2^ = 0.33 [0.28-0.43] (p < 0.001, Figure 7A) indicating modest predictability. In contrast, a model using high-throughput trafficking data alone produced an R^2^ = 0.65 [0.59-0.70] (p < 0.001, data not shown). Combining both the *in silico* and trafficking features to predict peak tail current density did not improve models using the trafficking feature alone (R^2^ = 0.65 [0.61-0.71]). Interestingly, this predictive performance was achieved though the trafficking data were derived using a homozygous expression system. In Figure 7A and B, we highlight three outliers significantly underestimated using *in silico* methods alone, Q84P, A79P, and L87P. These residues reside on an α-helix and are relatively tolerant to most non-proline substitutions (Figure 4). Proline substitutions are unique in the α-helical context as they do not have a free amide proton to allow for backbone hydrogen bonding and thus are disruptive to secondary structure.

**Figure 7:**
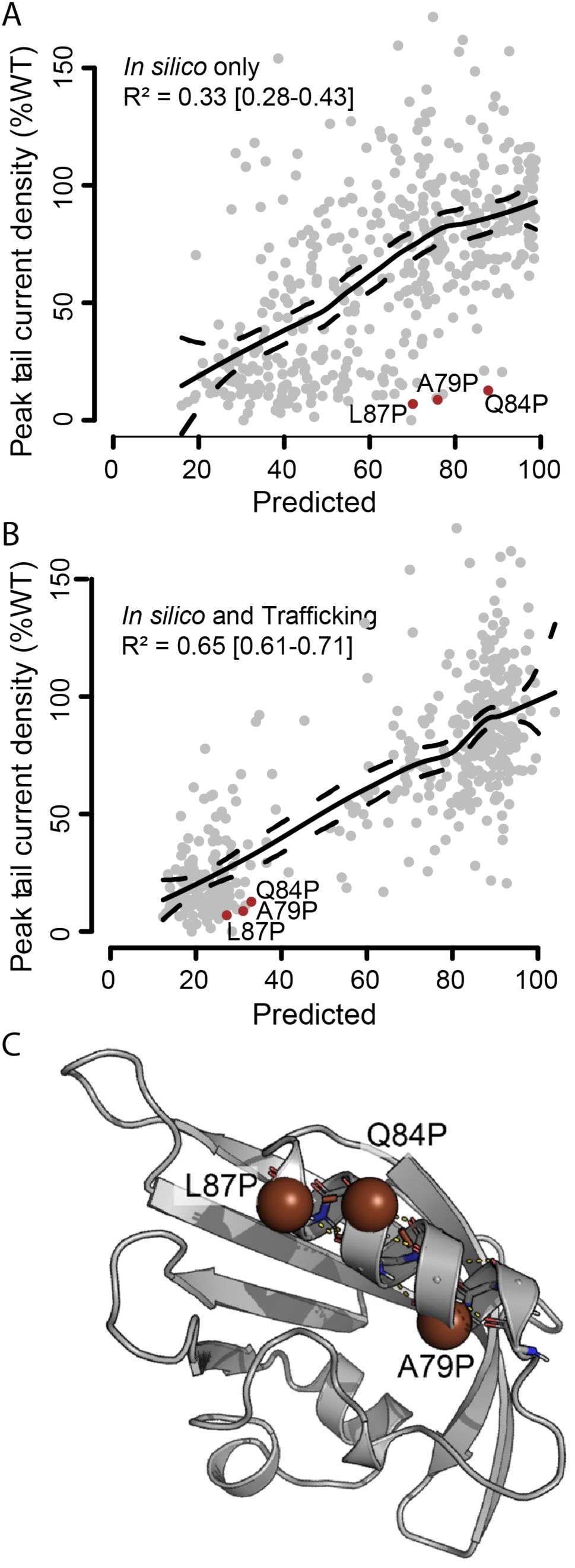
Modelling peak tail current density using trafficking and/or in silico variant classifiers. (A) Models of peak tail current density using *in silico* variant predictors REVEL, CardioBoost, and peak tail current density as predictive features (see Materials and Methods). Three variants with z-scores below two standard deviations of the mean are A79P, Q84P, and L87P are highlighted. (B) A regression of *in silico* and trafficking data to peak current density data. (C) These residues are on the c-terminal side of the longest α-helix in the PAS domain.

We additionally evaluated classification models using Area under the Receiver Operating Characteristic curves (AUC) to determine how well each feature could classify variants with loss of peak tail current density (Figure 8). REVEL and CardioBoost were nearly identical across a range of peak tail current density cut-offs of pathogenicity (AUC ∼ 0.80). Our structure-derived feature also correlated modestly with peak tail current density (AUC ∼ 0.75), though this feature added unique information to the classification of loss-of-function variants (as seen by higher AUCs when combined with REVEL and CardioBoost; Figure 8). Most notably, the trafficking data, evaluated at all thresholds of peak tail current density, substantially outperformed all other features combined or individually (Figure 8). Trafficking could correctly classify loss of peak tail current density variants with an AUC reaching 0.94; adding all *in silico* predictors to the trafficking data resulted in only a modest improvement with an AUC = 0.96.

**Figure 8:**
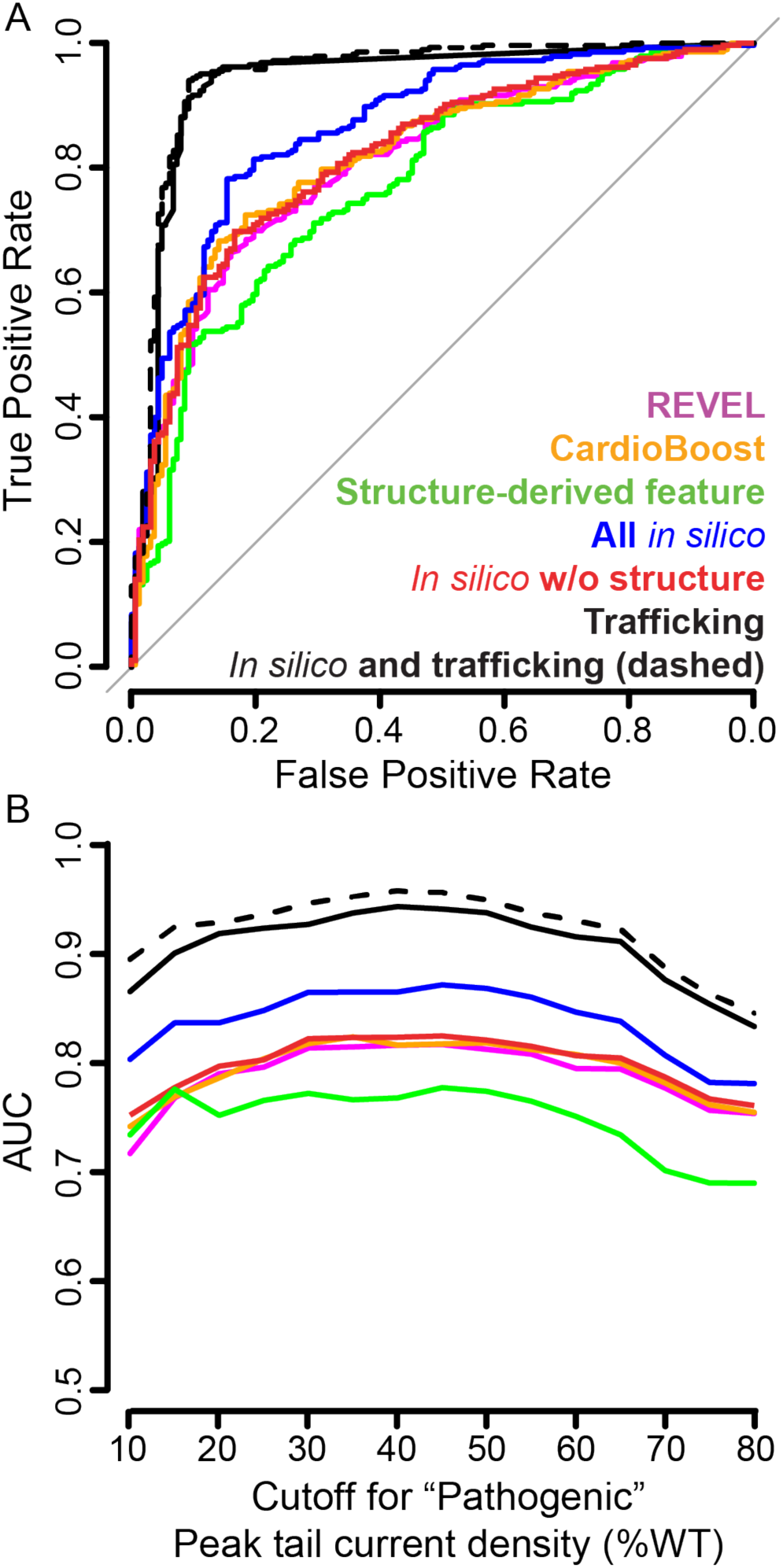
Receiving operating characteristic (ROC) curves predicting loss of peak tail current density and corresponding areas under the ROC curve (AUC). (A) ROC curves predicting loss of more than 50 % of WT peak tail current density for the exon 2 missense SNV. (B) AUCs calculated at multiple cut-offs of “pathogenicity”, or loss of peak tail current density, both above and below 50 % of WT.

## Discussion

Here, we compared and validated a massively parallel trafficking assay against a high information content automated patch clamp assay for assessing *KCNH2* variants. Although we found approximately 40 % of missense SNVs had loss-of-function (< 50 % of WT), approximately 55 % of variants resembled WT (Figures 1 and 2). Most variants that caused loss of function were dominant negative (Figure 5) and did not significantly alter gating properties of the channel (Figure 3). We also observed that deleterious variants were typically found together in clusters within the PAS domain (Figure 4). Finally, peak tail current density phenotype was very well predicted by the trafficking data while *in silico* variant tools provided little additional information to these predictions (Figures 7 and 8).

Like earlier studies (Kozek et al., 2020; Ng et al., 2020), we observed significant variability in the extent of loss of function amongst the variants characterized (Figures 1C and 2C). A structure-based analysis of the loss-of-function shows that loss-of-function variants were distributed throughout the PAS domain, though denser in certain pockets (Figure 4C). As expected, buried regions were generally intolerant of substitutions whilst solvent exposed regions were generally tolerant. For example, the helix stretching from Q75 to A89 has one face relatively intolerant to substitution pointed inward towards the core of the PAS domain while the remaining residues are exposed to solvent and more permissive to substitutions. Similary, I82 and L86 are oriented towards the core of the domain and variants at these sites have a mean trafficking score of 22 and 33 % of WT, respectively whilst R77, A80, Q84, and G88 are predominantly exposed to the cytosol and are correspondingly more tolerant to substitutions with a mean trafficking score of 95, 79, 68 and 83 % of WT (across all missense variants). The exception to this trend is proline substitutions: In the 15 residues which compose an α-helical connector mentioned above, there are 8/13 proline SNVs that induce >75 % loss-of-function. This is consistent with the proline substitution eliminating the backbone amid hydrogen necessary for hydrogen bonding in the α-helix; the subsequent destabilization is reflected in the loss of protein at the cell surface.

An advantage of patch clamp assays is the collection of gating and ion permeation as well as trafficking. The PAS domain is known to play a role in regulating channel deactivation (Gianulis & Trudeau, 2011; Ke et al., 2013). However, <5 % (21/458) SNV showed enhanced rates of deactivation and only 1 % (4/458) exhibited slow deactivation. Most PAS domain variants that affect channel deactivation also induce a moderate to severe trafficking defect (Ke et al., 2013). The severe trafficking defective variants (e.g. I31S, I42N, Y43C) do not pass the current density threshold needed for accurate analysis of gating phenotypes in our assay. For variants reported to have a modest trafficking defect and modest acceleration of channel deactivation (∼35-70 % of WT, K28E, F29L, N33T & R56Q (Gianulis & Trudeau, 2011)), our data is consistent with that in the literature. However, there are some variants (e.g. C64Y, T65P and I96T) that displayed a small perturbation in deactivation as homozygous variants (Ke et al., 2013) but are found to be WT-like in our assay. This discrepancy is likely due to the combined expression of the WT allele reducing the impact of the mild deactivation gating defect. A fortunate consequence of there being relatively few variants where gating defects are the major cause of loss-of-function is that the massively parallel trafficking assays can therefore correctly identify most loss-of-function variants.

Prospectively classifying variants has the potential to drastically reduce the delay between acquiring a genome sequencing result and establishing a diagnosis. Although bioinformatic tools are capable of prospectively characterizing variants, these tools are consistently too sensitive (Miosge et al., 2015). This high sensitivity is one reason the ACMG suggests *in silico* data be interpreted as supporting evidence (BP4/PP3) of variant pathogenicity compared to strong evidence (BS3/PS3) for functional data (Richards et al., 2015). For *KCNH2*, we observed around 55 % of SNVs were predicted pathogenic by REVEL and CardioBoost compared to around 40 % in our functional assays. Using these tools to classify peak tail current density-defective variants, this difference results in an AUC of ∼ 0.8 whereas trafficking classifies with an AUC > 0.9 (Figure 8). When directly modelling peak tail current density, *in silico* features combined could only explain 33 % of the variability, R^2^ = 0.33 [0.28-0.43]. High-throughput trafficking data alone produced an R^2^ = 0.65 [0.59-0.70]. These results indicate the trafficking data contain information that is not present in the *in silico* predictors evaluated. However, it remains to be seen whether the *in silico* features may still offer orthogonal information to the functional data when predicting the clinical significance of variants in *KCNH2*.

## Limitations

Although these results represent an exciting development in ion channel physiology and the assessment of variants in ion channelopathies, there are some potential limitations to the data and methods described. The assays described here were developed in HEK293 cells which may not express all the regulatory proteins or molecules that might be required for correct signalling or channel regulation. For this reason, there has been considerable interest in using induced pluripotent stem cell derived cardiac myocytes for modelling genetic heart diseases (Brandao et al., 2020; Ge et al., 2019). This limitation is less likely to be a major problem for *KCNH2* as these channels are not significantly modulated by signalling pathways (Vandenberg et al., 2012). In addition, gain-of-function variants in *KCNH2* channels have been associated with a distinct clinical phenotype, short QT syndrome (Gaita et al., 2003). The assays described are capable of detecting these gain-of-function variants, however, further development of this capability is part of ongoing work.

## Conclusions

To overcome the barrier of implementing genome-guided medicine, there is currently an unmet need for large-scale functional characterization of genetic variants. These results can be acquired prospectively and the data available for clinical geneticists contemporaneously with whole genome sequencing results. Massively parallel assays have the potential to produce high information content at scale and prospectively to reduce the number of VUSs. Here, we demonstrated how a trafficking assay of *KCNH2* could recapitulate peak tail current density, the benchmark assay for ion channel variant classification. These functional assays produce information not currently present in *in silico* predictors evaluated and therefore offer additional insight into variant pathogenesis. Furthermore, it is very likely this approach will be applicable to many other ion channels and membrane proteins where protein misfolding of variant proteins is the major cause of loss of function (Yue et al., 2005).

## Materials and Methods

Detailed methods are also available in a previous publication (Kozek et al., 2020). *Mutagenesis.* To manage *KCNH2* into workable sections, a tile system was created using the Quikchange Lightning Multi kit (Agilent), as reported earlier (Kozek et al., 2020). Specifically, the PAS domain was included in the first of these “tiles”, which we will refer to as the “PAS tile”, flanked by restriction sites for ClaI and SpeI. To enable cell surface labelling of K_V_11.1, an HA tag (NSEHYPYDVPDYAVTFE) was inserted in the region between amino acids T443 and E444, which was previously found to have no effect on the electrical or trafficking properties of the channel (Ficker et al., 2003).

We completed a comprehensive site saturation mutagenesis on the PAS tile with mutagenic primer pairs comprising a 5’NNN segment on the forward primer, where N is a mix of A/C/G/T. The intent was to generate a pool of primers that would generate a pool of plasmids with a relatively equal assortment of possible codon combinations. We purchased 231 PCR primers for the first 231 amino acids of K_V_11.1 we intended to mutate. These mutagenized plasmids were then constructed as previously described (Kozek et al., 2020). To investigate the success of the procedure and to quantify the quality and diversity of plasmid uptake, we grew colonies after electroporation in parallel with colony expansion in 50 mL of LB broth. Colonies were grown on LB-ampicillin agar plates at a dilution of 1:10,000. We recorded the total number of colonies produced, which informed the number of unique variant-barcoded plasmids.

### Linking *KCNH2* variants to the barcodes that reside on a plasmid

To be able to identify which barcodes belong to which variants, we needed to bring the variants closer to the barcodes to be within the range of current next generation sequencing technology. Using i7 and i5 primer pairs with specific adapter sequences and PCR, we generated multiple pools of sequences containing the barcodes and variant sequences in close proximity. Results were obtained in a fastq format and analyzed using Python and R studio. In this way, we paired barcodes and variants across the entire PAS tile.

### Transfecting and creating a stable cell line and preparing cells for flow sorting followed the same protocol as previously described (Kozek et al., 2020)

#### NovaSeq sequencing of the generated libraries

DNA was isolated from each of the H, M, L and Neg sorted pools using 100 μL QuickExtract (Lucigen) per 1 million cells, following the manufacturer’s instructions. PCR reaction, containing 1 μL of isolated DNA with 25-35 cycles, was carried out to amplify the barcode for Illumina sequencing using Q5 polymerase (NEB), according to the manufacturer’s instructions. These libraries were purified with AmpureXP beads (Beckman Coulter), following manufacturer’s instructions. The libraries were assessed with stringent quality control parameters and sequenced on a NovaSeq 6000 instrument with 150 base paired end sequencing. The resulting fastq files were analyzed using python and R studio. Variant counts from each pool of sorted cells were aggregated to calculate a trafficking score using the following equation:

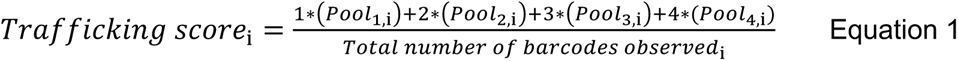

Where Pool_n,i_ is the fraction of the i^th^ barcode in the n^th^ pool of sorted cells; n ranges from 1 (no K_V_11.1 present, i.e. no AF647 signal) to 4 (high abundance of K_V_11.1, i.e. high AF647 signal). The scores were then aggregated by variant and normalized with a linear transformation so that trafficking score ranged from 0 (barcodes only observed in the AF647 negative pool) to 100 for WT. The scores were then averaged across the 2 replicate experiments (separate transfections of the mutant library pool).

#### Modeling peak tail current

Models included REVEL (Ioannidis et al., 2016), CardioBoost (Zhang et al., 2021), structurally derived parameter density (in three-dimensional space), and high-throughput trafficking data. Each feature was regressed using a restricted cubic spline (rcs from rms package in R (<u>rmsOverview function - RDocumentation</u>)) with four knots. Linear models were constructed using the lm function in the stats package in R (<u>lm function - RDocumentation</u>). Coefficients of Determination were adjusted for the number of regression features. Logistic regression models to evaluate ROC/AUCs were built using the glm function in the rms package in R.

#### Replications

Two mutagenized plasmids with unique barcode associations were generated as biological replicates. Each of these plasmids was transfected into three separate HEK cell lines for a total of six replicates. Within each sorted sample, there were approximately 2-9 (interquartile range) technical replicates owing to the redundancy of barcodes and codon substitutions associated with the same missense variant.

#### Automated patch clamp

A detailed method paper describing the design of heterozygous *KCNH2* vector, generation of Flp-In T-rex HEK293 *KCNH2* variant cell lines, cell culture routine of heterozygous KCNH2 Flp-In HEK293 for automated patch clamp electrophysiology, operation of SyncroPatch 384PE automated patch clamp, and voltage protocols and data analysis was recently published (Ng et al., 2021). Briefly, all 458 *KCNH2* variants used in the automated patch clamp were co-expressed with WT *KCNH2* from the same plasmid in stable Flp-In HEK293 cell lines. The expression of *KCNH2* in these Flp-In HEK293 cell lines was induced with doxycycline for 48 hours prior to undertaking automated patch clamp experiments. WT and a negative control were included together with 10 *KCNH2* variants in every SyncroPatch assay plate. Technical replicates were performed for all plates. Voltage protocols were designed to measure steady-state activation, onset of inactivation and steady-state deactivation (Ng et al., 2021). In total, 92 assay plates were used to collect >35,000 patch clamp experiments. All recordings were subjected to the following quality control measures (i.e. seal resistance > 300 M*Ω*, capacitance between 5 and 50 pF and leak corrected current measured at −120 mV leak step is within ± 40 pA of the baseline) to remove poor recordings before data analysis. Peak tail current was quantified by measuring the peak amplitude of the tail current at −120 mV, after a depolarizing step to +40 mV for 1 s. Current values were normalised to cell capacitance to obtain current density (pA/pF). Steady-state activation was derived from a 1 s isochronal activation protocol, cells were depolarized to potentials from –50 mV to +70 mV in 10 mV increments for 1 s before stepping to –120 mV to record tail currents. A Boltzmann equation was fitted to the −120 mV peak tail currents to determine the mid-point of the voltage dependence of activation (V_0.5_):

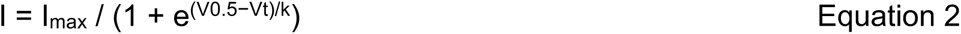

where: V_0.5_ is the half-maximum activation voltage, V_t_ is the test potential, k is the slope factor and I_max_ is the maximum tail current. To assess deactivation, cells were depolarized to +40 mV for 1 s before repolarization to potentials in the range +20 to −150 mV in 10 mV decrements for 3 s. The tail current represents the recovery from inactivation followed by channel deactivation. The extent of channel deactivation was directly quantified by measuring the peak current at −50 mV and 500 ms later, which is reported as the current decay ratio at 500 ms. The time constant of deactivation was also determined by fitting the decaying portion of the tail current recordings using a double exponential function. The overall time constant of deactivation was calculated as a weighted sum of the two components using Equation 3:

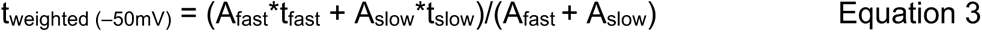

where A is the current amplitude and t is the time constant for the fast and slow components. Channel inactivation was assessed by measuring the ratio of the peak tail current amplitudes at −50 mV and −120 mV recorded from the deactivation protocol above. The rate of onset of inactivation was measured using a triple pulse protocol where cells were depolarised to +40 mV for 1 s to fully activate and then inactivate the channels before stepping to −110 mV for 10 ms to allow channels to recover from inactivation into the open state. Membrane potential was then depolarised to voltages in the range +60 to −20 mV (in 10 mV decrements) to inactivate channels. A single exponential was fitted to the inactivating current trace to determine the time constant for the onset of inactivation. To minimise the impact of poor voltage clamp control during large outward current traces we only fitted the exponential function to the current trace after it had fallen to <500 pA, which resulted in fewer n numbers for this analysis.

In addition, if the n number that pass quality control for analysis of gating parameters is less than 25 % of the recordings that have passed quality control parameters for peak tail current density then these variants are classified as not analysed (N/A) for the relevant gating parameters in the result.

## Acknowledgements

This work was funded by a NSW Cardiovascular Disease Senior Scientist Grant (JIV), an National Health and Medical Research Council Principal Research Fellowship (JIV), National Institutes of Health grant R00HL135442 (BMK), Leducq Foundation for Cardiovascular Research grant 18CVD05 “Towards Precision Medicine with Human iPSCs for Cardiac Channelopathies” (BMK). We also acknowledge support of the Victor Chang Cardiac Research Institute Innovation Centre, funded by the NSW Government.

## Author Contribution statement

CAN undertook patch clamp assays, analyzed data, prepared figures and wrote manuscript. RU undertook trafficking assays, analyzed the data and wrote manuscript. JF developed automated analysis routines and assisted with analysis of patch clamp data. APH assisted with development of automated patch clamp protocols, analysis routines and helped draft paper. KAK analyzed data and drafted manuscript. LRV undertook trafficking assays. DM undertook trafficking assays. BMK conceived project, assisted with data analysis and interpretation, wrote manuscript. JIV conceived project, assisted with data analysis and interpretation, wrote manuscript.

---

**Supplementary Figure 1:**
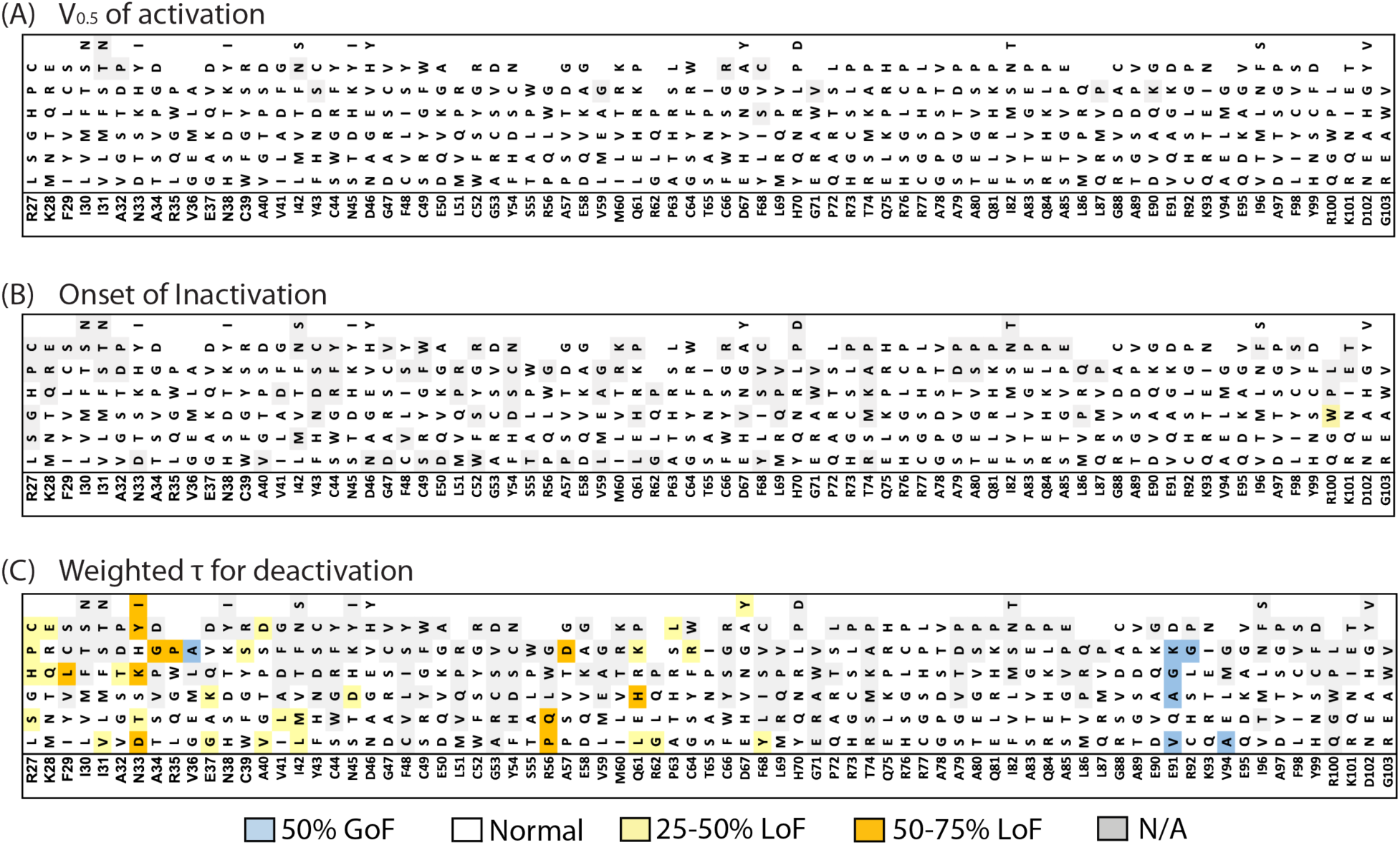
Onset of channel inactivation (top) and weighted *τ* measured at –50 mV for channel deactivation. When the n numbers that pass quality control for analysis of gating parameters is less than 25 % of the recordings that have passed quality control parameters for peak tail current density then these variants are classified as not analyzed (N/A).

**Table S1:**
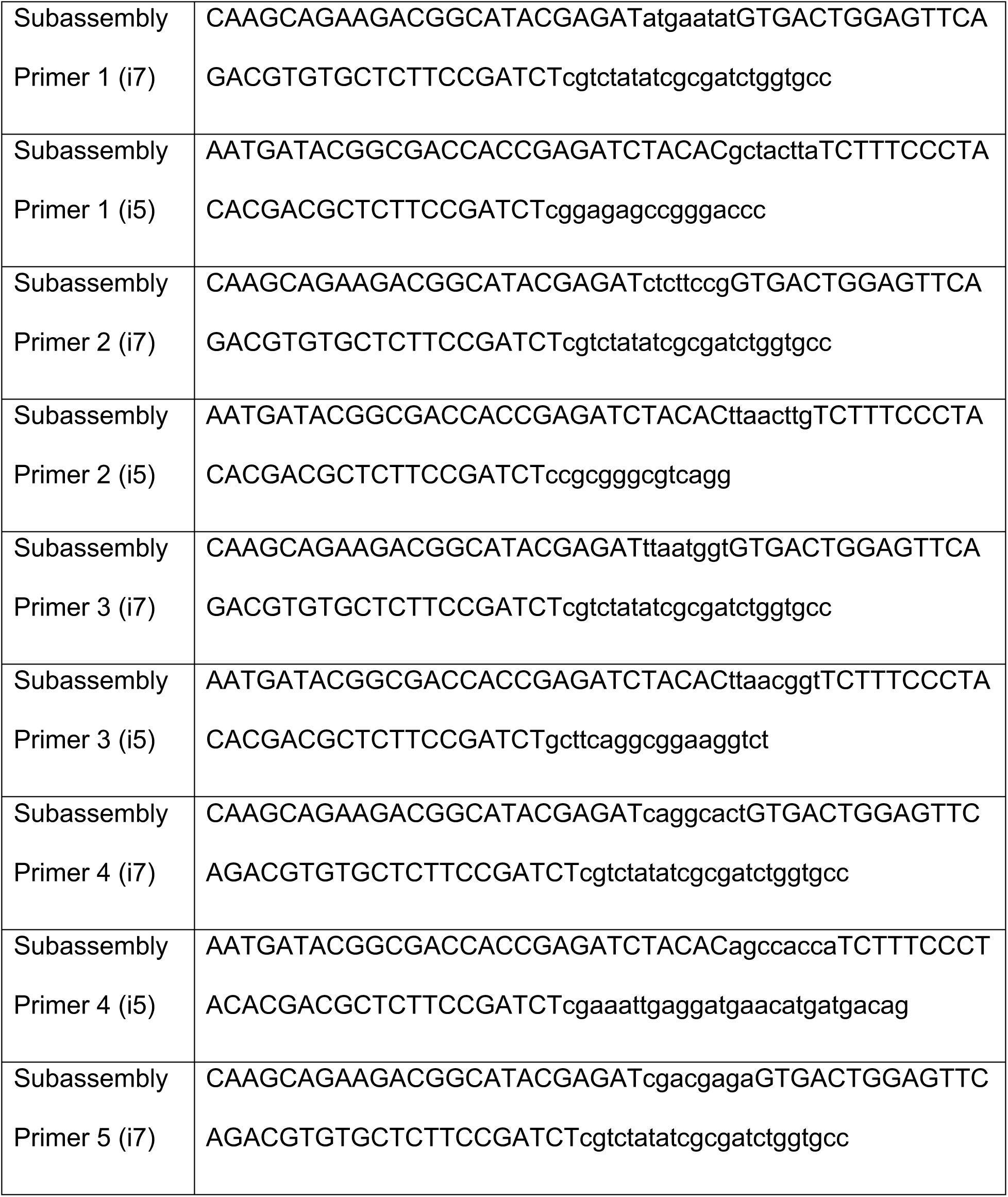
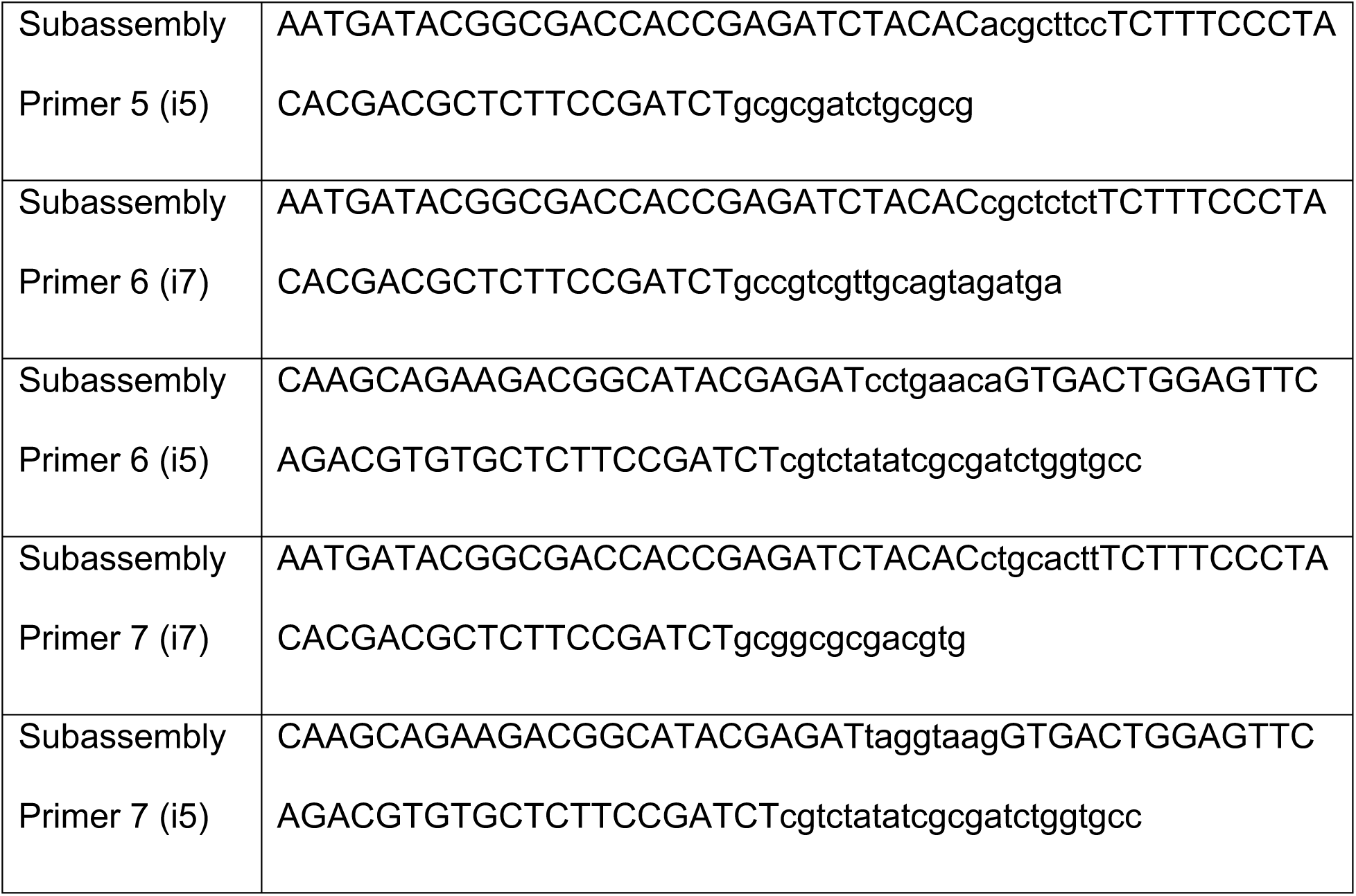
Primers pairs (i7 and i5 indices) used to link barcodes to variants.

**Table S2:**
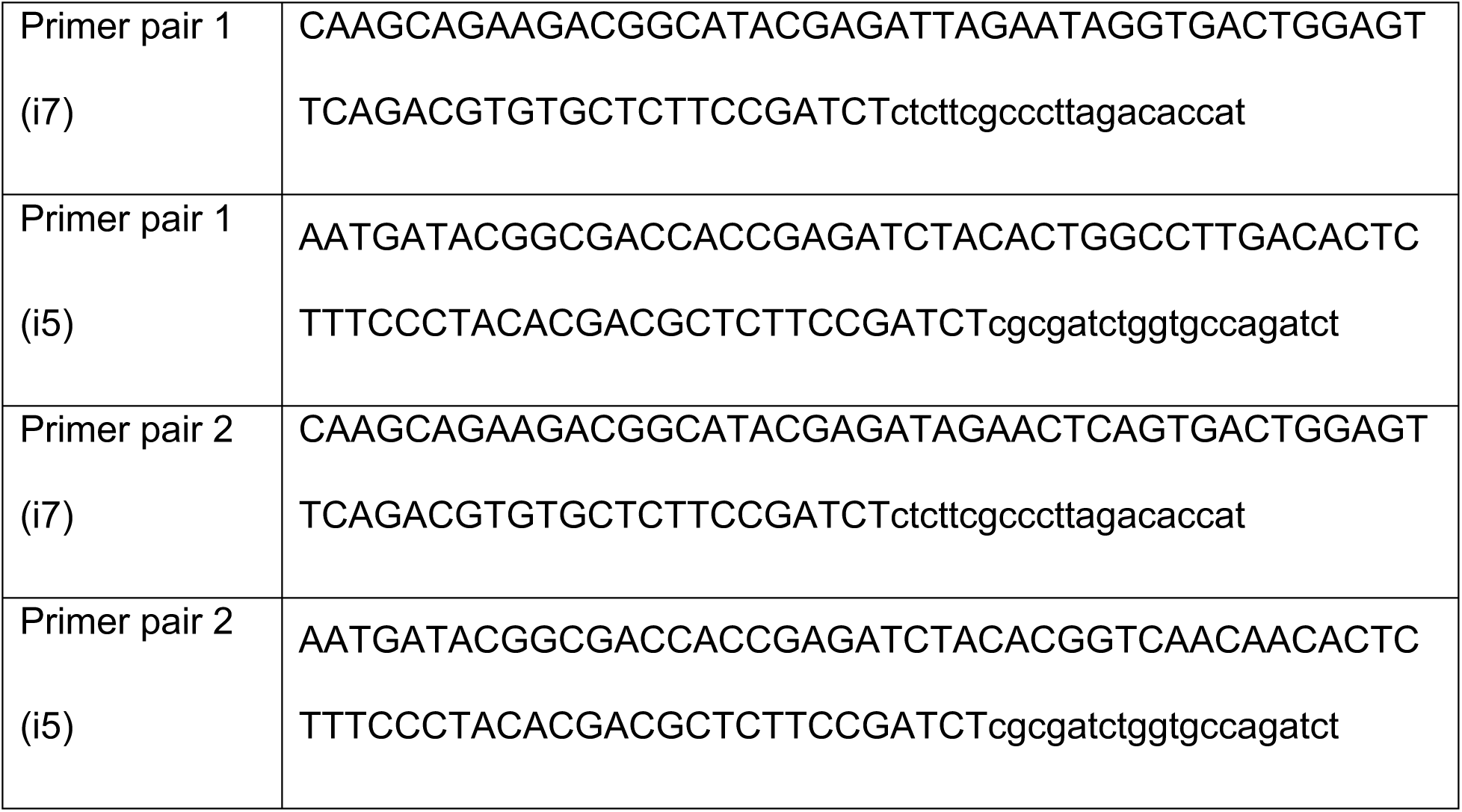
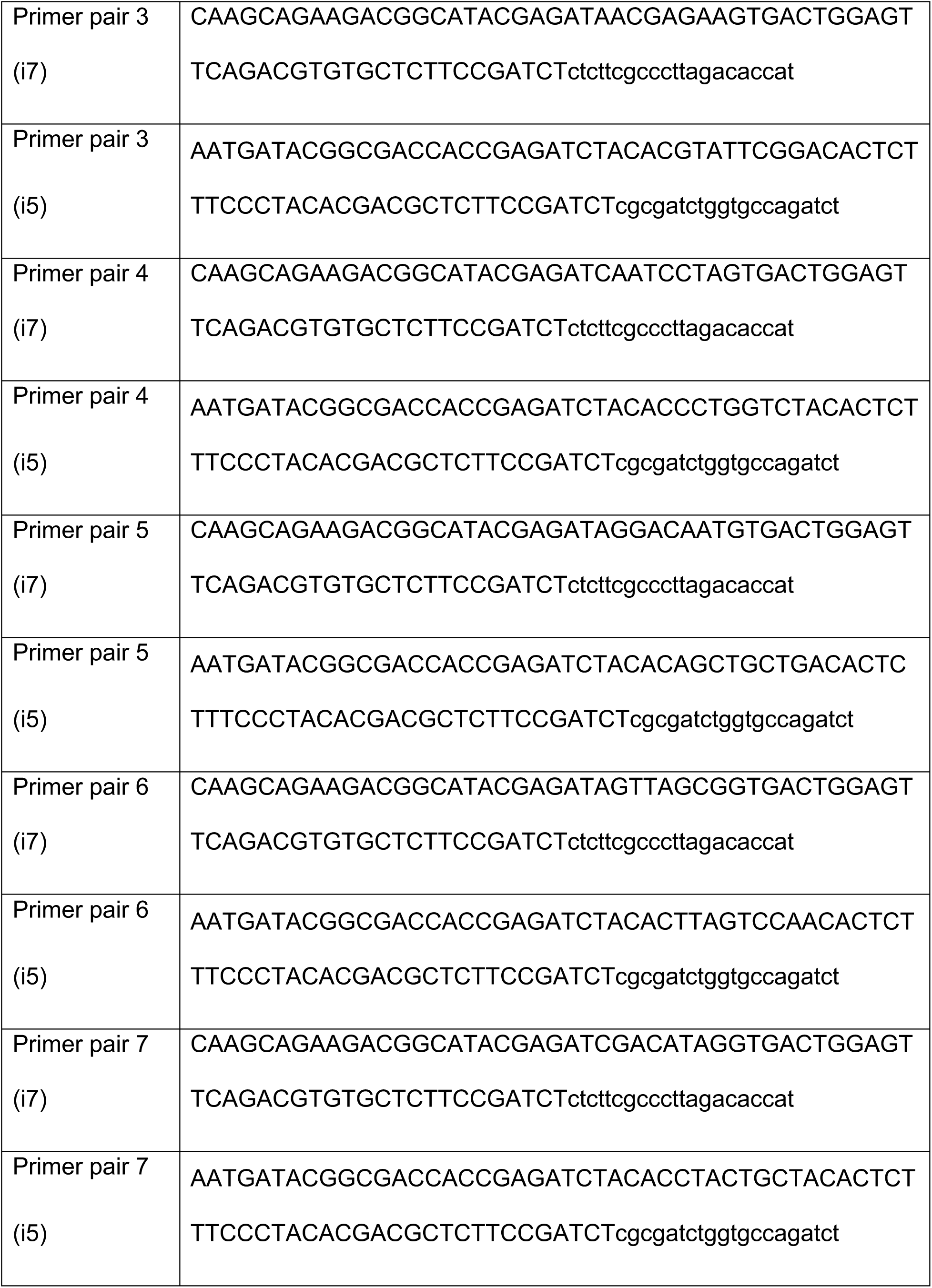
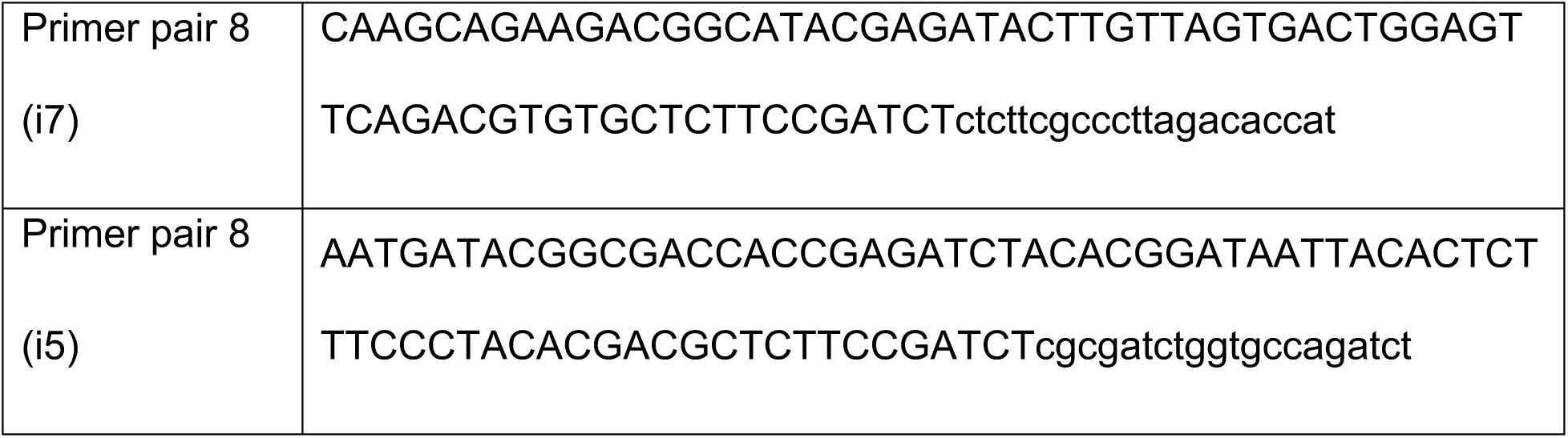
Primer pairs used for analysing variant trafficking in the HEK cells.

## References

Anderson, C. L., Kuzmicki, C. E., Childs, R. R., Hintz, C. J., Delisle, B. P., & January, C. T. (2014). Large-scale mutational analysis of Kv11.1 reveals molecular insights into type 2 long QT syndrome [Research Support, N.I.H., Extramural Research Support, Non-U.S. Gov’t]. Nat Commun, 5, 5535. https://doi.org/10.1038/ncomms6535

Brandao, K. O., van den Brink, L., Miller, D. C., Grandela, C., van Meer, B. J., Mol, M. P. H., de Korte, T., Tertoolen, L. G. J., Mummery, C. L., Sala, L., Verkerk, A. O., & Davis, R. P. (2020, Nov 10). Isogenic Sets of hiPSC-CMs Harboring Distinct KCNH2 Mutations Differ Functionally and in Susceptibility to Drug-Induced Arrhythmias. Stem Cell Reports, 15(5), 1127–1139. https://doi.org/10.1016/j.stemcr.2020.10.005

Delisle, B. P., Anson, B. D., Rajamani, S., & January, C. T. (2004, Jun 11). Biology of cardiac arrhythmias: ion channel protein trafficking. Circulation Research, 94(11), 1418–1428. https://doi.org/10.1161/01.RES.0000128561.28701.ea

Ficker, E., Dennis, A. T., Wang, L., & Brown, A. M. (2003, Jun 27). Role of the cytosolic chaperones Hsp70 and Hsp90 in maturation of the cardiac potassium channel HERG. Circulation Research, 92(12), e87–100. https://doi.org/10.1161/01.RES.0000079028.31393.15

Findlay, G. M., Daza, R. M., Martin, B., Zhang, M. D., Leith, A. P., Gasperini, M., Janizek, J. D., Huang, X., Starita, L. M., & Shendure, J. (2018, Oct). Accurate classification of BRCA1 variants with saturation genome editing. Nature, 562(7726), 217–222. https://doi.org/10.1038/s41586-018-0461-z

Gaita, F., Giustetto, C., Bianchi, F., Wolpert, C., Schimpf, R., Riccardi, R., Grossi, S., Richiardi, E., & Borggrefe, M. (2003, Aug 26). Short QT Syndrome: a familial cause of sudden death. Circulation, 108(8), 965–970. https://doi.org/10.1161/01.CIR.0000085071.28695.C4

Ge, N., Liu, M., Ding, Y., Krawczyk, J., McInerney, V., Galvin, J., Shen, S., Prendiville, T., & O’Brien, T. (2019, Dec). Generation and characterization of twelve human induced pluripotent stem cell (iPSC) lines from four familial long QT syndrome type 1 (LQT1) patients carrying KCNQ1 c.1201dupC mutation. Stem Cell Res, 41, 101650. https://doi.org/10.1016/j.scr.2019.101650

Gianulis, E. C., & Trudeau, M. C. (2011, Jun 24). Rescue of aberrant gating by a genetically encoded PAS (Per-Arnt-Sim) domain in several long QT syndrome mutant human ether-a-go-go-related gene potassium channels [Research Support, N.I.H., Extramural Research Support, Non-U.S. Gov’t]. Journal of Biological Chemistry, 286(25), 22160–22169. https://doi.org/10.1074/jbc.M110.205948

Glazer, A. M., Kroncke, B. M., Matreyek, K. A., Yang, T., Wada, Y., Shields, T., Salem, J. E., Fowler, D. M., & Roden, D. M. (2020, Feb). Deep Mutational Scan of an SCN5A Voltage Sensor. Circ Genom Precis Med, 13(1), e002786. https://doi.org/10.1161/CIRCGEN.119.002786

Glazer, A. M., Wada, Y., Li, B., Muhammad, A., Kalash, O. R., O’Neill, M. J., Shields, T., Hall, L., Short, L., Blair, M. A., Kroncke, B. M., Capra, J. A., & Roden, D. M. (2020, Jul 2). High-Throughput Reclassification of SCN5A Variants. American Journal of Human Genetics, 107(1), 111–123. https://doi.org/10.1016/j.ajhg.2020.05.015

Ioannidis, N. M., Rothstein, J. H., Pejaver, V., Middha, S., McDonnell, S. K., Baheti, S., Musolf, A., Li, Q., Holzinger, E., Karyadi, D., Cannon-Albright, L. A., Teerlink, C. C., Stanford, J. L., Isaacs, W. B., Xu, J., Cooney, K. A., Lange, E. M., Schleutker, J., Carpten, J. D., Powell, I. J., Cussenot, O., Cancel-Tassin, G., Giles, G. G., MacInnis, R. J., Maier, C., Hsieh, C. L., Wiklund, F., Catalona, W. J., Foulkes, W. D., Mandal, D., Eeles, R. A., Kote-Jarai, Z., Bustamante, C. D., Schaid, D. J., Hastie, T., Ostrander, E. A., Bailey-Wilson, J. E., Radivojac, P., Thibodeau, S. N., Whittemore, A. S., & Sieh, W. (2016, Oct 6). REVEL: An Ensemble Method for Predicting the Pathogenicity of Rare Missense Variants. American Journal of Human Genetics, 99(4), 877–885. https://doi.org/10.1016/j.ajhg.2016.08.016

January, C. T., Gong, Q., & Zhou, Z. (2000, Dec). Long QT syndrome: cellular basis and arrhythmia mechanism in LQT2. Journal of Cardiovascular Electrophysiology, 11(12), 1413–1418. https://doi.org/10.1046/j.1540-8167.2000.01413.x

Kalia, S. S., Adelman, K., Bale, S. J., Chung, W. K., Eng, C., Evans, J. P., Herman, G. E., Hufnagel, S. B., Klein, T. E., Korf, B. R., McKelvey, K. D., Ormond, K. E., Richards, C. S., Vlangos, C. N., Watson, M., Martin, C. L., & Miller, D. T. (2017, Feb). Recommendations for reporting of secondary findings in clinical exome and genome sequencing, 2016 update (ACMG SF v2.0): a policy statement of the American College of Medical Genetics and Genomics. Genet Med, 19(2), 249–255. https://doi.org/10.1038/gim.2016.190

Ke, Y., Ng, C. A., Hunter, M. J., Mann, S. A., Heide, J., Hill, A. P., & Vandenberg, J. I. (2013, Aug 15). Trafficking defects in PAS domain mutant Kv11.1 channels: roles of reduced domain stability and altered domain-domain interactions [Research Support, Non-U.S. Gov’t]. Biochemical Journal, 454(1), 69–77. https://doi.org/10.1042/BJ20130328

Keating, M. T., & Sanguinetti, M. C. (2001, Feb 23). Molecular and cellular mechanisms of cardiac arrhythmias. Cell, 104(4), 569–580. https://doi.org/10.1016/s0092-8674(01)00243-4

Kozek, K. A., Glazer, A. M., Ng, C. A., Blackwell, D., Egly, C. L., Vanags, L. R., Blair, M., Mitchell, D., Matreyek, K. A., Fowler, D. M., Knollmann, B. C., Vandenberg, J. I., Roden, D. M., & Kroncke, B. M. (2020, Dec). High-throughput discovery of trafficking-deficient variants in the cardiac potassium channel KV11.1. Heart Rhythm, 17(12), 2180–2189. https://doi.org/10.1016/j.hrthm.2020.05.041

Kroncke, B. M., Glazer, A. M., Smith, D. K., Blume, J. D., & Roden, D. M. (2018, May). SCN5A (NaV1.5) Variant Functional Perturbation and Clinical Presentation: Variants of a Certain Significance. Circ Genom Precis Med, 11(5), e002095. https://doi.org/10.1161/CIRCGEN.118.002095

Kroncke, B. M., Mendenhall, J., Smith, D. K., Sanders, C. R., Capra, J. A., George, A. L., Blume, J. D., Meiler, J., & Roden, D. M. (2019). Protein structure aids predicting functional perturbation of missense variants in SCN5A and KCNQ1. Comput Struct Biotechnol J, 17, 206–214. https://doi.org/10.1016/j.csbj.2019.01.008

Landrum, M. J., Lee, J. M., Benson, M., Brown, G. R., Chao, C., Chitipiralla, S., Gu, B., Hart, J., Hoffman, D., Jang, W., Karapetyan, K., Katz, K., Liu, C., Maddipatla, Z., Malheiro, A., McDaniel, K., Ovetsky, M., Riley, G., Zhou, G., Holmes, J. B., Kattman, B. L., & Maglott, D. R. (2018, Jan 4). ClinVar: improving access to variant interpretations and supporting evidence. Nucleic Acids Res, 46(D1), D1062–D1067. https://doi.org/10.1093/nar/gkx1153

Lek, M., Karczewski, K. J., Minikel, E. V., Samocha, K. E., Banks, E., Fennell, T., O’Donnell-Luria, A. H., Ware, J. S., Hill, A. J., Cummings, B. B., Tukiainen, T., Birnbaum, D. P., Kosmicki, J. A., Duncan, L. E., Estrada, K., Zhao, F., Zou, J., Pierce-Hoffman, E., Berghout, J., Cooper, D. N., Deflaux, N., DePristo, M., Do, R., Flannick, J., Fromer, M., Gauthier, L., Goldstein, J., Gupta, N., Howrigan, D., Kiezun, A., Kurki, M. I., Moonshine, A. L., Natarajan, P., Orozco, L., Peloso, G. M., Poplin, R., Rivas, M. A., Ruano-Rubio, V., Rose, S. A., Ruderfer, D. M., Shakir, K., Stenson, P. D., Stevens, C., Thomas, B. P., Tiao, G., Tusie-Luna, M. T., Weisburd, B., Won, H. H., Yu, D., Altshuler, D. M., Ardissino, D., Boehnke, M., Danesh, J., Donnelly, S., Elosua, R., Florez, J. C., Gabriel, S. B., Getz, G., Glatt, S. J., Hultman, C. M., Kathiresan, S., Laakso, M., McCarroll, S., McCarthy, M. I., McGovern, D., McPherson, R., Neale, B. M., Palotie, A., Purcell, S. M., Saleheen, D., Scharf, J. M., Sklar, P., Sullivan, P. F., Tuomilehto, J., Tsuang, M. T., Watkins, H. C., Wilson, J. G., Daly, M. J., MacArthur, D. G., & Exome Aggregation, C. (2016, Aug 18). Analysis of protein-coding genetic variation in 60,706 humans. Nature, 536(7616), 285–291. https://doi.org/10.1038/nature19057

Miosge, L. A., Field, M. A., Sontani, Y., Cho, V., Johnson, S., Palkova, A., Balakishnan, B., Liang, R., Zhang, Y., Lyon, S., Beutler, B., Whittle, B., Bertram, E. M., Enders, A., Goodnow, C. C., & Andrews, T. D. (2015, Sep 15). Comparison of predicted and actual consequences of missense mutations. Proceedings of the National Academy of Sciences of the United States of America, 112(37), E5189–5198. https://doi.org/10.1073/pnas.1511585112

Morais Cabral, J. H., Lee, A., Cohen, S., Chait, B., Li, M., & Mackinnon, R. (1998). Crystal structure and functional analysis of the HERG potassium channel N terminus: a eukaryotic PAS domain. Cell, 95, 649–655. http://www.ncbi.nlm.nih.gov/entrez/query.fcgi?db=pubmed&cmd=Retrieve&dopt=AbstractPlus&list_uids=5432560159351622838related:tswXl65UZEsJ

Ng, C. A., Farr, J., Young, P., Windley, M. J., Perry, M. D., Hill, A. P., & Vandenberg, J. I. (2021). Heterozygous KCNH2 variant phenotyping using Flp-In HEK293 and high-throughput automated patch clamp electrophysiology. Biol Methods Protoc, 6(1), bpab003. https://doi.org/10.1093/biomethods/bpab003

Ng, C. A., Perry, M. D., Liang, W., Smith, N. J., Foo, B., Shrier, A., Lukacs, G. L., Hill, A. P., & Vandenberg, J. I. (2020, Mar). High-throughput phenotyping of heteromeric human ether-a-go-go-related gene potassium channel variants can discriminate pathogenic from rare benign variants. Heart Rhythm, 17(3), 492–500. https://doi.org/10.1016/j.hrthm.2019.09.020

Priori, S. G., Wilde, A. A., Horie, M., Cho, Y., Behr, E. R., Berul, C., Blom, N., Brugada, J., Chiang, C. E., Huikuri, H., Kannankeril, P., Krahn, A., Leenhardt, A., Moss, A., Schwartz, P. J., Shimizu, W., Tomaselli, G., & Tracy, C. (2013, Dec). HRS/EHRA/APHRS expert consensus statement on the diagnosis and management of patients with inherited primary arrhythmia syndromes: document endorsed by HRS, EHRA, and APHRS in May 2013 and by ACCF, AHA, PACES, and AEPC in June 2013. Heart Rhythm, 10(12), 1932–1963. https://doi.org/10.1016/j.hrthm.2013.05.014

Rajamani, S., Anderson, C. L., Anson, B. D., & January, C. T. (2002, Jun 18). Pharmacological rescue of human K(+) channel long-QT2 mutations: human ether-a-go-go-related gene rescue without block. Circulation, 105(24), 2830–2835. https://doi.org/10.1161/01.cir.0000019513.50928.74

Richards, S., Aziz, N., Bale, S., Bick, D., Das, S., Gastier-Foster, J., Grody, W. W., Hegde, M., Lyon, E., Spector, E., Voelkerding, K., Rehm, H. L., & Committee, A. L. Q. A. (2015, May). Standards and guidelines for the interpretation of sequence variants: a joint consensus recommendation of the American College of Medical Genetics and Genomics and the Association for Molecular Pathology. Genet Med, 17(5), 405–424. https://doi.org/10.1038/gim.2015.30

Roden, D. M., & Balser, J. R. (1999, Nov). A plethora of mechanisms in the HERG-related long QT syndrome. Genetics meets electrophysiology. Cardiovascular Research, 44(2), 242–246. https://doi.org/10.1016/s0008-6363(99)00224-2

Semsarian, C., Ingles, J., Ross, S. B., Dunwoodie, S. L., Bagnall, R. D., & Kovacic, J. C. (2021, May 25). Precision Medicine in Cardiovascular Disease: Genetics and Impact on Phenotypes: JACC Focus Seminar 1/5. Journal of the American College of Cardiology, 77(20), 2517–2530. https://doi.org/10.1016/j.jacc.2020.12.071

Shimizu, W., Moss, A. J., Wilde, A. A., Towbin, J. A., Ackerman, M. J., January, C. T., Tester, D. J., Zareba, W., Robinson, J. L., Qi, M., Vincent, G. M., Kaufman, E. S., Hofman, N., Noda, T., Kamakura, S., Miyamoto, Y., Shah, S., Amin, V., Goldenberg, I., Andrews, M. L., & McNitt, S. (2009, Nov 24). Genotype-phenotype aspects of type 2 long QT syndrome. Journal of the American College of Cardiology, 54(22), 2052–2062. https://doi.org/10.1016/j.jacc.2009.08.028

Starita, L. M., Ahituv, N., Dunham, M. J., Kitzman, J. O., Roth, F. P., Seelig, G., Shendure, J., & Fowler, D. M. (2017, Sep 7). Variant Interpretation: Functional Assays to the Rescue. American Journal of Human Genetics, 101(3), 315–325. https://doi.org/10.1016/j.ajhg.2017.07.014

Vandenberg, J. I., Perry, M. D., Perrin, M. J., Mann, S. A., Ke, Y., & Hill, A. P. (2012, Jul). hERG K(+) channels: structure, function, and clinical significance [Research Support, Non-U.S. Gov’t Review]. Physiological Reviews, 92(3), 1393–1478. http://www.ncbi.nlm.nih.gov/pubmed/22988594

Vanoye, C. G., Desai, R. R., Fabre, K. L., Gallagher, S. L., Potet, F., DeKeyser, J. M., Macaya, D., Meiler, J., Sanders, C. R., & George, A. L., Jr. (2018, Nov). High-Throughput Functional Evaluation of KCNQ1 Decrypts Variants of Unknown Significance. Circ Genom Precis Med, 11(11), e002345. https://doi.org/10.1161/CIRCGEN.118.002345

Yue, P., Li, Z., & Moult, J. (2005, Oct 21). Loss of protein structure stability as a major causative factor in monogenic disease. Journal of Molecular Biology, 353(2), 459–473. https://doi.org/10.1016/j.jmb.2005.08.020

Zhang, X., Walsh, R., Whiffin, N., Buchan, R., Midwinter, W., Wilk, A., Govind, R., Li, N., Ahmad, M., Mazzarotto, F., Roberts, A., Theotokis, P. I., Mazaika, E., Allouba, M., de Marvao, A., Pua, C. J., Day, S. M., Ashley, E., Colan, S. D., Michels, M., Pereira, A. C., Jacoby, D., Ho, C. Y., Olivotto, I., Gunnarsson, G. T., Jefferies, J. L., Semsarian, C., Ingles, J., O’Regan, D. P., Aguib, Y., Yacoub, M. H., Cook, S. A., Barton, P. J. R., Bottolo, L., & Ware, J. S. (2021, Jan). Disease-specific variant pathogenicity prediction significantly improves variant interpretation in inherited cardiac conditions. Genet Med, 23(1), 69–79. https://doi.org/10.1038/s41436-020-00972-3

